# Fine tuning the mechanism of Antigen 43 self-association to modulate aggregation levels of *Escherichia coli* pathogens

**DOI:** 10.1101/2021.04.19.440493

**Authors:** Julieanne L Vo, Gabriela C Martínez Ortiz, Makrina Totsika, Alvin Lo, Andrew E. Whitten, Lilian Hor, Kate M Peters, Valentin Ageorges, Nelly Caccia, Mickaël Desvaux, Mark A Schembri, Jason J Paxman, Begoña Heras

**Affiliations:** Department of Biochemistry and Genetics, La Trobe Institute for Molecular Science, La Trobe University, Melbourne VIC 3086, Australia; Institute of Health and Biomedical Innovation, Centre for Immunology and Infection Control, School of Biomedical Sciences, Queensland University of Technology, Herston, QLD 4006, Australia; Australian Infectious Diseases Research Centre, School of Chemistry and Molecular Biosciences, The University of Queensland, Brisbane QLD 4072, Australia; Australian Centre for Neutron Scattering, Australian Nuclear Science and Technology Organisation, Lucas Heights, NSW 2234, Australia; Université Clermont Auvergne, INRAE, UMR454 MEDiS, 63000, Clermont-Ferrand, France

**Author notes:** These authors contributed equally: Julieanne L Vo, Gabriela C Martínez Ortiz. These authors jointly supervised this work: Jason J Paxman, Begoña Heras. Correspondence and requests for materials should be addressed to J.J.P. or to B.H.

## Abstract

Bacterial aggregates and biofilms allow bacteria to colonise a diverse array of surfaces that can ultimately lead to infections, where the protection they afford permits bacteria to resist anti-microbials and host immune factors. Despite these advantages there is a trade-off, whereby bacterial spread is reduced. As such, biofilm development needs to be regulated appropriately to suit the required niche. Here we investigate members from one of largest groups of bacterial adhesins, the autotransporters, for their critical role in the formation of bacterial aggregates and biofilms. We describe the structural and functional characterisation of autotransporter Ag43 homologues from diverse pathogenic *Escherichia* strains. We reveal a common mode of trans-association that leads to cell clumping and show that subtle variations in these interactions governs their aggregation kinetics. Our in depth investigation reveals an underlying molecular basis for the ‘tuning’ of bacterial aggregation.

## Introduction

Many bacterial species have the ability to transition from a free-swimming lifestyle to the formation of aggregated or biofilm communities (1). These multicellular structures offer several adaptation advantages, such as enhanced resistance against antimicrobial agents, chemical detergents and immune factors (2) and account for 65 to 80% of bacterial infections in humans (3). Although very common and often competitively beneficial to bacteria, the molecular mechanisms underlying bacterial self-recognition and aggregation remain to be fully elucidated. For example, there is increasing evidence that bacterial aggregation can be coordinated by controlled chemotactic motility (4–7) as well as entropic forces that drive the depletion of substances which block cell-cell interactions (8). In relation to this study, bacterial aggregation is also known to be facilitated by proteins that decorate the cell surface (2), but the precise motifs and mechanisms of self-recognition remain largely uncharacterised.

A class of proteins often associated with the bacterial aggregation phenotype is the autotransporter (AT) superfamily (2, 9). These proteins belong to the largest group of secreted and outer-membrane proteins in Gram-negative bacteria, and share a common translocation pathway defined by the type V secretion system (9–11). Within this family, the self-associating autotransporters (SAATs), are principally involved in bacterial aggregation and biofilm formation (12).

The best-characterised SAAT, antigen 43 (Ag43), is an *Escherichia coli* protein that drives cell aggregation and biofilm formation (9, 13–15). Ag43 possesses a modular domain organisation that is common throughout the autotransporters, which encompasses an N-terminal signal sequence that directs the secretion of the protein across the inner membrane, a passenger (α) domain that is responsible for the function of the protein, and a C-terminal translocator (β) domain that facilitates the translocation of the functional α-domain to the cell surface (16–19). After translocation, the Ag43 α-domain (α^43^) is cleaved but remains attached to the cell via non-covalent interactions (9, 13, 20-23).

Ag43 exhibits allelic variation based on sequence diversity, particularly within its functional α-domain, and this can lead to altered degrees of aggregation (15, 20). A recent comprehensive analysis of Ag43 functional α-domains identified four major phylogenetic groups (C1-C4), which differed in amino acid sequence as well as auto-aggregative phenotypes (20). In the case of the Ag43a variant, structural analysis has revealed it possesses an L-shaped beta-helical conformation that forms a ‘head-to-tail’ self-association interaction between neighbouring cells to drive bacterial aggregation (13). Importantly, this work identified two patches of charged/polar residues that constitute self-association interfaces within its α-domain (α^43a^) (13). However, whether this head-to-tail mechanism of self-association is conserved across all Ag43 variants and AT in general, and how sequence changes impact these interactions, remain unanswered questions.

In this study, we have determined the crystal structures of a set of Ag43 homologues from three pathogenic *E. coli* strains; the enterohemorrhagic *E. coli* (EHEC) O157:H7 strain EDL933 and the uropathogenic *E. coli* (UPEC) strains UTI89 and CFT073. These prototype *E. coli* pathogens cause significant human infections including haemorrhagic colitis and haemolytic-uremic syndrome by EHEC (24–26) and urinary tract infections and sepsis by UPEC (27). Importantly, together they encode Ag43 homologues that represent its phylogenetic diversity. Our data demonstrates that the functional α-domain of these Ag43 proteins folds into a bent three-stranded β-helix, which is predicted to be a structural hallmark of the SAAT family. Through structural and functional studies, we have identified the residues on the surface of each Ag43 α-domain that mediate trans-association with other α^43^ molecules, which leads to bacterial clumping. Our results also uncovered subtle differences in α^43^-α^43^ oligomerisation modes, which determine distinct aggregation kinetics and interaction strengths. Overall, we demonstrate a common mode of self-association among Ag43 proteins that drives head-to-tail associations that can be fine-tuned to attain diverse aggregation activities. We propose this may represent a universal mechanism of aggregation across the large family of SAATs and may contribute to colonisation of different ecological niches.

## Results

### Ag43 homologues promote differences in bacterial aggregation

We first examined the aggregation properties of a set of Ag43 homologues from different *E. coli* pathogens that represent the phylogenetic diversity in the Ag43 family (20). Specifically, we focused on Ag43 from EHEC EDL933 (Ag43^EDL933^) and UPEC UTI89 (Ag43^UTI89^), which belong to the C2 and C4 Ag43 phylogenetic classes, respectively (20, 28, 29). We also selected Ag43b from UPEC CFT073, which belongs to the C3 group (14, 20); Ag43a, which we have characterised previously and also belongs to the C3 group (13, 20), served as a control. Each gene was PCR amplified and cloned into the expression vector pBAD/Myc-His A, the resultant plasmids were transformed into the *E. coli fim agn43* null strain MS528 (30), and their ability to promote bacterial aggregation was compared using a sedimentation assay that measured the settling kinetics of standing cultures (Fig. 1A). Recombinant MS528 strains harbouring all of the different Ag43 variants were observed to auto-aggregate. However, there were clear differences in the aggregation kinetics; while Ag43^EDL933^ and Ag43a sedimented rapidly, Ag43^UTI89^ and Ag43b exhibited a slower sedimentation profile (Fig. 1A). This difference was most apparent in the early stage of bacterial aggregation (30 mins), with fluorescence microscopy of agn43-negative gfp-positive *E. coli* K-12 strain OS56 harbouring the different Ag43 variants, revealing larger aggregates for Ag43^EDL933^ and Ag43a compared to Ag43^UTI89^ and Ag43b (Fig. 1B). Using flow cytometry, we compared Ag43^EDL933^ to Ag43b on either side of the sedimentation range, again confirming the higher aggregation of Ag43^EDL933^ (Fig. 1C). As another measure of aggregation/biofilm kinetics we used the biofilm ring test, an assay whereby the movement of microbeads in microplates is blocked in the course of biofilm formation. In this assay we used the recombinant MS528 *E. coli* strain harbouring all of the different Ag43 variants. The most marked difference was observed after six hours of *E. coli* growth whereby both Ag43a and Ag43^EDL933^ expressing cells completely blocked the microbeads contrary to Ag43b and Ag43^UTI89^ (Fig. 1D). Hence, *E. coli* cells expressing Ag43a and Ag43^EDL933^ appeared to sediment more rapidly due to the formation of larger aggregates, which in turn resulted in more rapid biofilm formation than Ag43^UTI89^ and Ag43b.

**Fig. 1.**
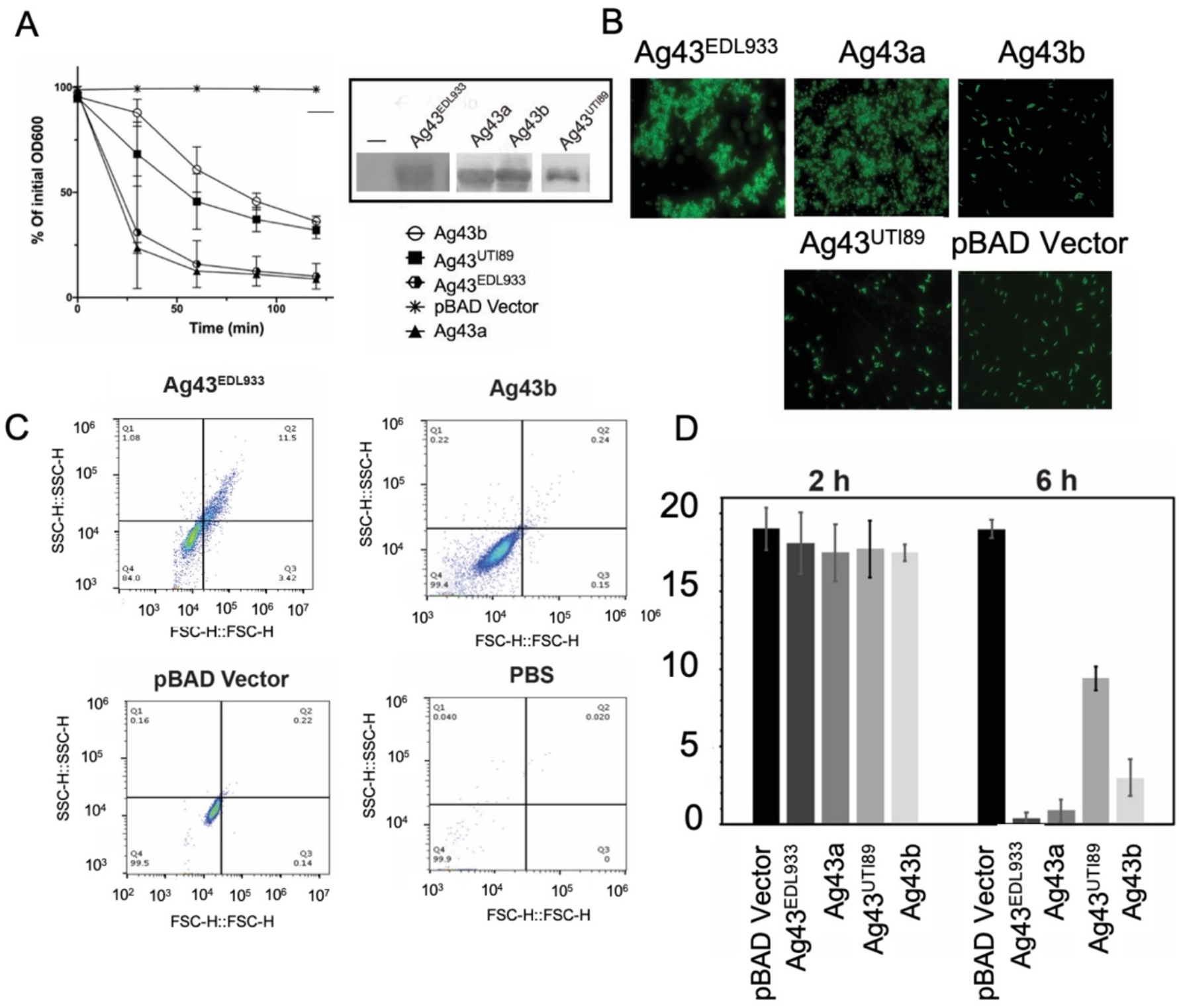
Aggregation kinetics of *E. coli* expressing the different Ag43 homologues. (*A*) *E. coli fim agn43* null strain MS528 expressing Ag43a, Ag43b, Ag43^UTI89^ and Ag43^EDL933^ from pBAD/Myc-His A were left to sediment for 2 hours with OD_600_ measurements taken at 30 min intervals. Experiments were performed in triplicate using pBAD/Myc-His A as a negative control. Ag43 protein production at the bacterial cell surface was examined using western blot analysis. (*B*) Representative images of GFP-positive *E. coli* K-12 (strain OS56) harbouring the same Ag43 plasmids were acquired using a Zeiss Axioplan 2 fluorescence microscope, at an early 30 min timepoint. (*C*) Forward vs side scatter plot for flow cytometry performed on *E. coli* K-12 strain OS56 harbouring pBAD/Myc-His A Ag43^EDL933^, Ag43b, and plasmid alone and PBS. *(D)* The biofilm ring test was performed on *E. coli fim agn43* null strain MS528 expressing Ag43a, Ag43b, Ag43^UTI89^ and Ag43^EDL933^ from pBAD/Myc-His A in microplates. Measurements are shown after 2 and 6 hours of growth. Results are expressed as biofilm formation indices (BFI), with the lower values indicating stronger biofilm formation.

### Diverse Ag43 homologues adopt L-shaped β-helical structures

The different functional properties of Ag43 homologues prompted us to characterise their atomic structures, noting that the α^43_EDL933^, α^43_UTI89^ and α^43b^ share 45%, 49% and 85% sequence identity, respectively, with α^43a^ (SI Appendix Table S1). The crystal structures of α^43_UTI89^ and α^43b^ were solved by molecular replacement using the previously determined structure of α^43a^ (PDB: 4KH3) (13) as a search model (Fig. 2). The structure of α^43_^EDL933 was solved using α^43_UTI89^ as a reference (Fig. 2). Crystals of α^43_UTI89^ and α^43b^ belong to the *C*2 and *P*2_1_22_1_ space groups, respectively, and each contained one molecule per asymmetric unit (Table 1). The structures of α^43_UTI89^ and α^43b^ were refined to 2.4 and 2.08 Å resolution and final R_factor_ values of 0.16 and 0.17 (R_free_ 0.21 and 0.20 respectively) (Table 1).

**Table 1.**
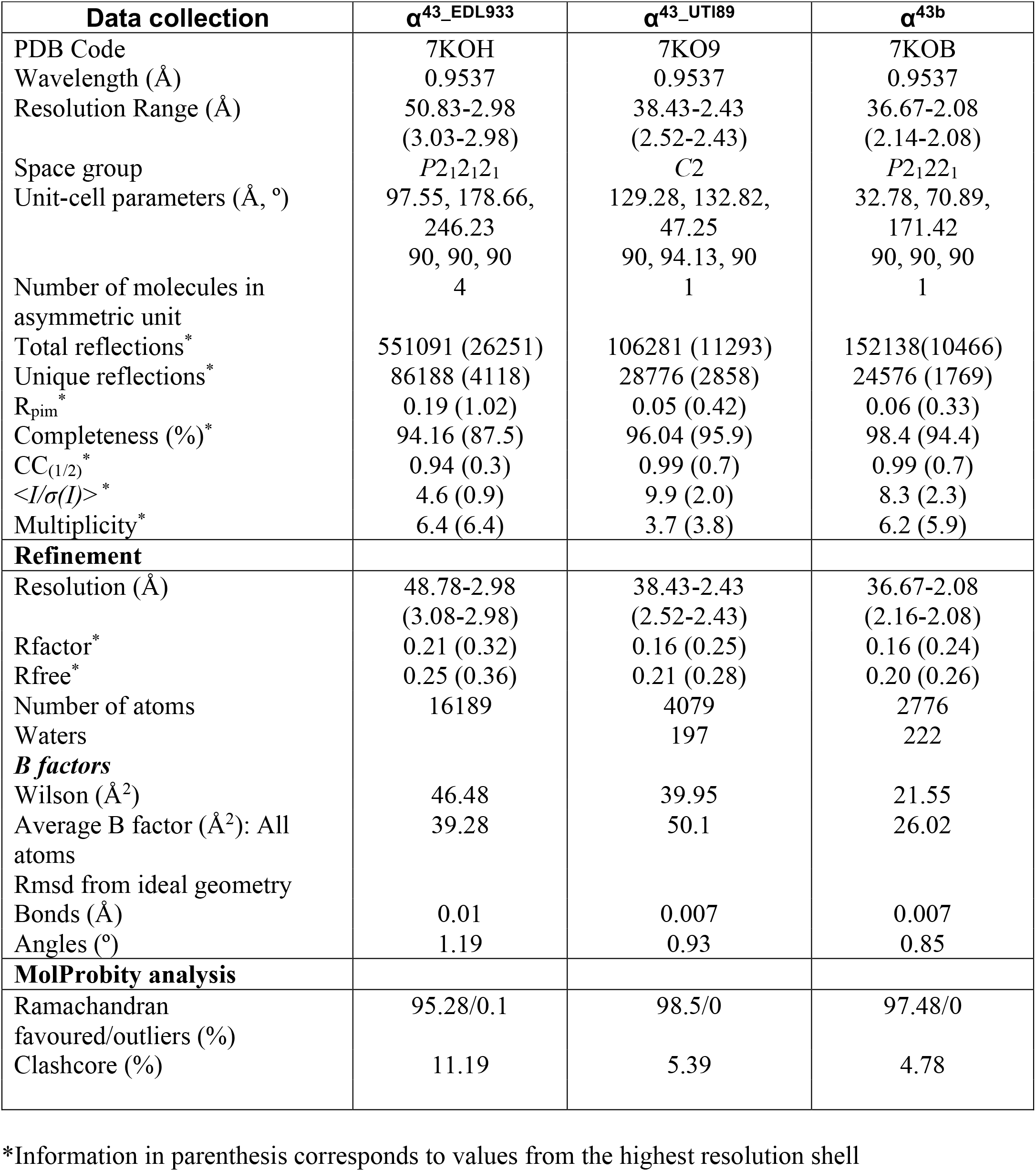
X-ray Crystallography data collection and statistics

**Fig. 2.**
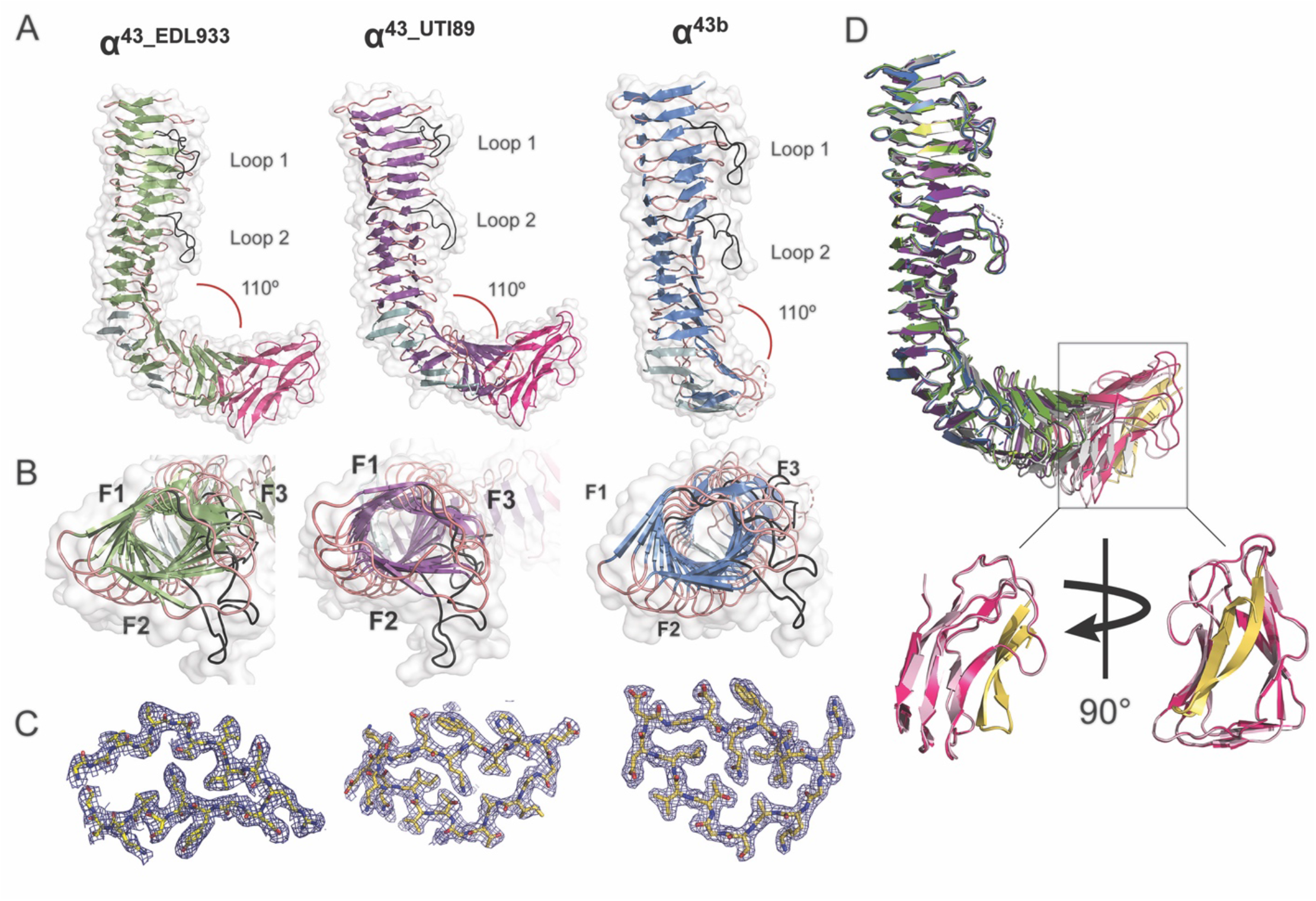
Structures of the α-domains of Ag43^EDL_933^ (α^43_EDL933^), Ag43^UTI89^ (α^43_UTI89^) and Ag43b (α^43b^). (*A*) *(B)* Side and top cartoon representations with β-strands in green, purple and blue, respectively, and turns are coloured in salmon. The two loops (loop 1 and loop 2) that protrude from the β-helices are shown in black, while the four β-hairpins appear in cyan. The AC domain of both α^43_EDL933^ and α^43_UTI89^ are coloured in hot pink. (*C*) Electron density map contoured at 1σ of a cross-section of α^43_EDL933^ (residues ^110^G-^129^G), α^43_UTI89^ (residues ^100^N-^120^K) and α^43b^ (residues ^90^L-^106^I). *(D)* Superimposition of α^43_EDL933^, α^43_UTI89^ and α^43b^ on that of α^43a^ with the latter depicted in grey. The AC domains are displayed in hot pink and light pink for α^43_EDL933^ and α^43_UTI89^ respectively, with the β-hairpin motifs yellow.

α^43_EDL933^ crystals belonged to the space group *P*2_1_2_1_2_1_, diffracting to 2.98 Å and contained four molecules per asymmetric unit. Following several steps of model building and refinement the structure of α^43_EDL933^ was refined to a R_factor_ of 0.21 (R_free_ of 0.25) (Table 1). Pairwise comparison of the four symmetrically independent monomers gave an r.m.s.d value for all Cα atoms of around 0.6 Å. As there were no significant differences observed between the molecules in the asymmetric unit, all subsequent structural comparisons were performed using the best-defined monomer.

Similar to α^43a^ (13), the α-domain of all three Ag43 homologues folds into an L-shaped right-handed three-stranded β-helix, with each turn of the β-spine comprising three β-strands linked by loop regions (Fig. 2A). In all cases, the long arm of the L-shape is formed by 13 turns, with the bent region formed by a pair of β-hairpin motifs on either side of three β-helix rungs, followed by a shorter C-terminal β-helical domain. α^43_UTI89^ and α^43_EDL933^ are shorter than α^43a^ and α^43b^; sequence alignment of these proteins showed that the former proteins are reduced in size by 69 amino acids in the α-domain (SI Appendix Fig. S1), which results in C-terminal β-helix domain consisting of four rungs instead of seven in α^43a^ (Fig. 2A). Although α^43a^ and α^43b^ are similar in size, the α^43b^ crystallised in this work is truncated and lacks the bottom section of the L-shaped β-helix (Fig. 2A). The α^43b^ truncation occurred during recombinant protein purification with cleavage occurring at residue 415 by an unknown protease. In all proteins, two loops protrude from the β-helix between rungs 2 and 3 (loop 1) and rungs 7 and 8 (loop 2). Both loop 1 and 2 consist on average of 15 residues, that include negatively charged residues which create acidic patches that protrude from the β-spine (SI Appendix Fig. S2). No functional role has been attributed to these protruding loops to date (13).

### α^43_UTI89^ and α^43_EDL933^ structures incorporate the autochaperone domain

During translocation to the bacterial cell surface, the functional α-domain of Ag43 adhesins is cleaved but remains attached to the C-terminal β-domain via non-covalent interactions (13, 17). The specific cleavage site varies among Ag43 homologues but typically localises between the β-helix and the predicted autochaperone (AC) domain (31). The α^43_UTI89^ and α^43_EDL933^ structures in this work correspond to the uncleaved α-domains.

Present within the structures of α^43_UTI89^ and α^43_EDL933^, their autochaperone regions consist of three β-strand rungs capped by a β-hairpin motif (Fig. 2D). This domain shows structural similarities with previously characterised autotransporter domains; where they superimpose with r.m.s.d values ranging between 1.50 - 4.37 Å with the AC domains from EspP (PDB: 3SZE), IcsA (PDB: 3ML3), p69 (PDB: 1DAB), Hap (PDB: 3SYJ) and Hbp (PDB: 1WXR) (32–37) (SI Appendix Table S1). Previously, the AC has been found to be involved in α-domain folding (38–41).

In the α^43_UTI89^ and α^43_EDL933^ structures the main β-helix and autochaperone domains are covalently linked via an extended 18 amino acid loop. Presumably due to flexibility, this loop in the α^43_UTI89^ (434–451) and α^43_EDL933^ (435–452) is mostly undefined in the electron density maps.

### Ag43^UTI89^ and Ag43b self-associate via an interface with fewer interactions

Ag43a from CFT073 drives bacterial aggregation by homotypic interactions between α^43a^ molecules from neighbouring cells in a head-to-tail conformation (13). To investigate whether this mechanism of self-association is conserved in Ag43^UTI89^, which shows a slow aggregating phenotype, we first examined its oligomerisation in the crystal lattice. Generation of crystallographically related molecules of α^43_UTI89^ revealed a head-to-tail dimer repeated along the crystal, which resembled the α^43a^-α^43a^ functional dimer (Fig. 4D) (13). However, while α^43a^ trans-dimers are stabilised by two interfaces consisting of a total of 18 hydrogen bonds and two salt bridges, α^43_UTI89^ dimers are mediated by a single interface with 13 hydrogen bonds [G13-N137, G30-N100, T32-N100 (two hydrogen bonds), N60-N100, N60-T98, D79-D79, T80-N60, T98-N60, T98-G30, N100-T32 (two hydrogen bonds), N137-G13] (Fig. 3A). To test if this α^43_UTI89^ crystallographic interface was indeed involved in mediating bacterial aggregation, we constructed a mutant with seven amino acid substitutions (Ag43^UTI89-*mt*^: T32G, N60G, D79G, T80G, T98G, N100G, N137G). Plasmid constructs harbouring full length Ag43^UTI89^ (pBAD::Ag43^UTI89^) or Ag43^UTI89-*mt*^ (pBAD::Ag43^UTI89-*mt*^), were transformed into the non-aggregative *E. coli* MG1655 *fim agn43* null strain (MS528) (42) and tested in cell-aggregation assays. Modification of the single interface in Ag43^UTI89^ completely abolished the ability of this protein to promote bacterial aggregation (Fig. 3B). This confirmed the self-association interface observed in the crystal lattice. This loss of function was not due to lack of expression of Ag43^UTI89^ on the bacterial surface as shown by western blot analysis of heat-released proteins (Fig. 3B, inset).

**Fig. 3.**
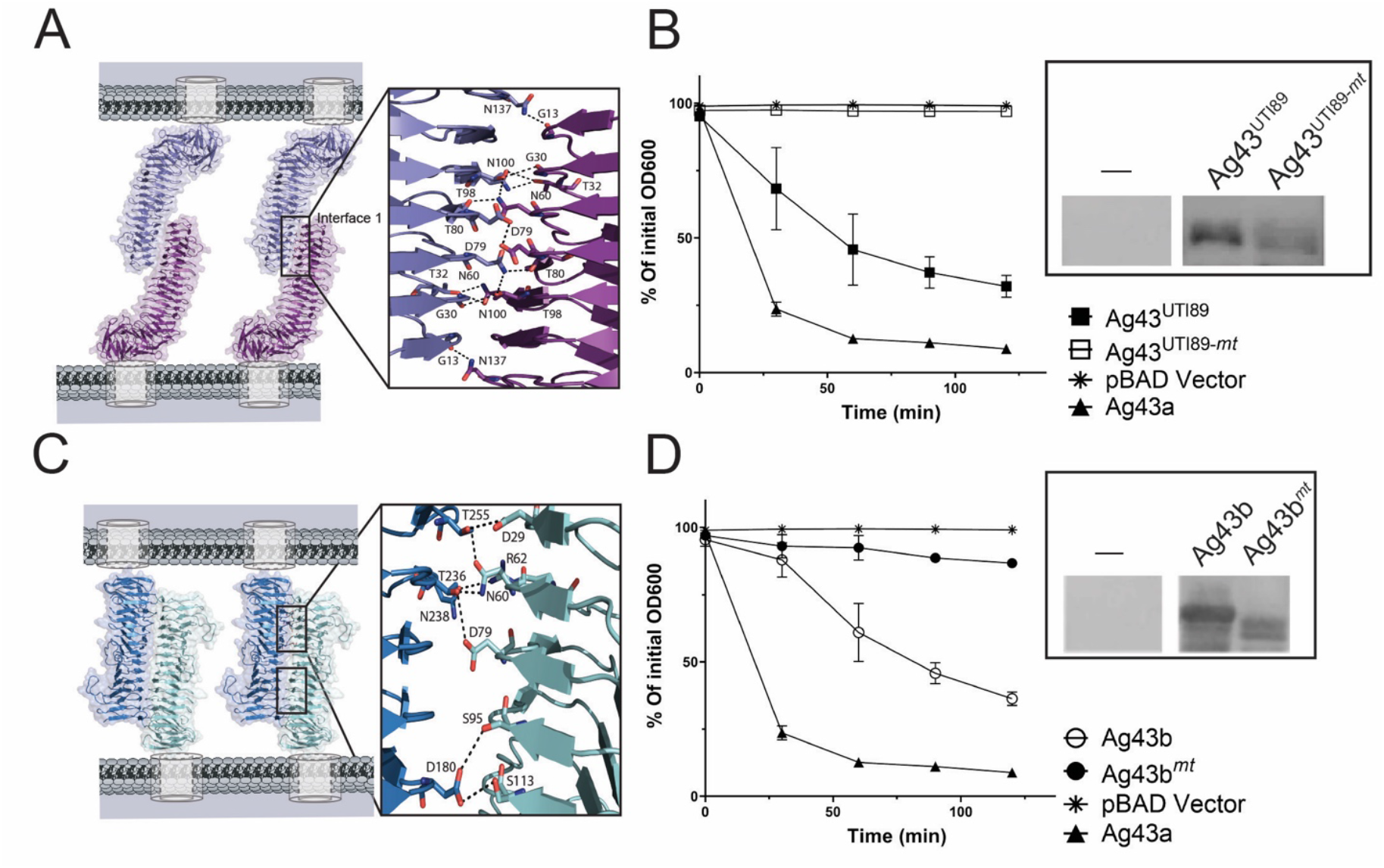
Ag43 self-association interfaces with fewer interactions. (*A*) Self-association of α^43_UTI89^ via a single interface consisting of thirteen hydrogen bonds [G13-N137, G30-T98, T32-N100 (two hydrogen bonds), N60-T98, N60-T80, D79-D79, T80-N60, T98-G30, T98-N60, N100-T32 (two hydrogen bonds) and N137-G13]. *(B)* Confirmation of the α^43_UTI89^ self-association interface shown by the aggregation profiles of *E. coli* strains expressing the interface mutant Ag43^UTI89-*mt*^ (T32G, N60G, D79G, T80G, T98G, N100G, N137G), WT Ag43^UTI89^, WT Ag43a and vector only pBAD/Myc-His A. The expression of Ag43 proteins on the bacterial cell surface was shown by western blot analysis (inset). (*C*) Self-association of α^43b^ via a double interface with each consisting of seven hydrogen bonds [D29-T255, N60-T255, R62-N238, D79-T236, S95-D180, S113-D180 and N60-T236]. *(D)* Confirmation of the α^43b^ self-association interface shown by the aggregation profiles of *E. coli* strains expressing the interface mutant Ag43b^*mt*^ (D29G, N60G, R62G, D79G and S95G), WT Ag43b, WT Ag43a and vector only pBAD/Myc-His A. The expression of Ag43 proteins on the bacterial cell surface was examined via western blot analysis (inset).

**Fig. 4.**
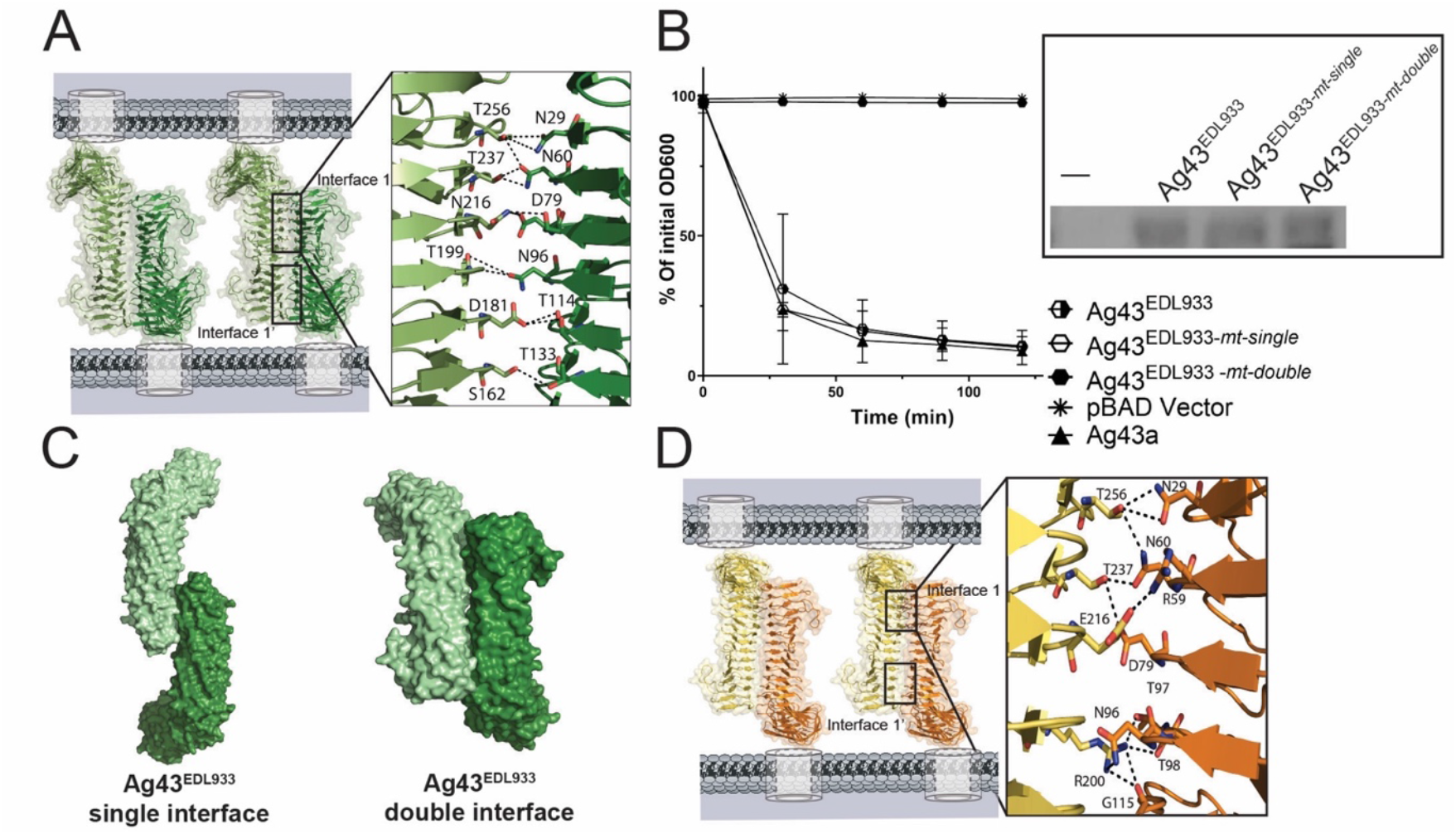
Ag43 interfaces with extensive self-association interfaces. (*A*) Double self-association interface of α^43_EDL933^, with each interface consisting of 12 hydrogen bonds [D181-T114 (2 H-bonds), T256-N29 (2 H-bonds), S162 –T133, N216-D79, N216-S78, T237-N60 (2 H-bonds) T256-N60, N96-T199 (2 H-bonds)]. (*B*) Confirmation of the α^43_EDL933^ double interface was shown by the loss of aggregation for the double interface mutant Ag43^EDL933-*mt-double*^ (T199G and T256G). In contrast, mutation of the predicted single interface Ag43^EDL933-*mt-single*^ (T15G, T32G, N60G, D79G, T98G, N100G, N119G and N138G) showed no effect on aggregation. Aggregation assays included WT Ag43^EDL933^, WT Ag43a and vector only pBAD/Myc-His A. The expression of Ag43 proteins on the bacterial cell surface was examined via western blot analysis (inset). (*C*) A depiction of the two alternative modes of α^43_EDL933^ self-association using a single or double interface. (*D*) Self-association of α^43a^ with each interface comprising 9 hydrogen bonds and a salt bridge [N29-T256 (2 H-bonds), N60-T256, N60-T237, D79-T237, N96-R200, T97-R200, T98-R200, G115-R200 and R59-E216 (salt bridge)] (13)

Although Ag43a and Ag43b share 85% sequence identity in their α-domains, these proteins differ significantly in their aggregation profiles (Fig. 1A) (14). Unlike α^43a^, α^43b^ did not crystallise in its dimeric form. However, given the similarity between α^43a^ and α^43b^, we superimposed two α^43b^ structures on the α^43a^-α^43a^ functional dimer (13) to infer the potential α^43b^ dimerisation interface (Fig. 3C). The predicted interaction surface consisted of two interfaces encompassing a total of 14 hydrogen bonds (D29-T255, N60-T255, N60-T236, R62-N238, D79-T236, S95-D180, S113-D180, in each interface). To confirm if this interface was indeed responsible for the aggregation phenotype of Ag43b, we mutated to glycine residues D29, N60, R62, D79 and S95 (Ag43b^*mt*^) and tested an MS528 strain expressing this mutant protein in aggregation assays. Substitutions of these five interface residues completely abolished the ability of Ag43b to mediate cell–cell aggregation compared with Ag43b WT, confirming the predicted mode of self-association (Fig. 3D).

### Ag43^EDL933^ self-associates through a double interface with extensive interactions

The α^43_EDL933^ crystals with four molecules in the asymmetric unit revealed complex non-crystallographic and crystallographic oligomers, however none of these resembled previously defined functional α^43^ dimers where the β-helical molecules coil around each other in trans-configuration (13) (SI Appendix Fig. S3). To define the self-association interface that leads to Ag43^EDL933^ bacterial clumping, α^43_EDL933^ molecules were superimposed onto α^43a^ dimers, stabilised by two interfaces (13) (Fig. 4C), and onto α^43_UTI89^ dimers with a single interacting interface (Fig. 4C; SI Appendix Fig. S4). All possible interactions at the interfaces of both models were predicted by exploring the different rotamer conformations of the residues mapping near the putative interfaces. The first superposition predicted an α^43_EDL933^ functional dimer stabilised by two interfaces, each consisting of 12 hydrogen bonds [D181-T114 (2 H-bonds), T256-N29 (2 H-bonds), S162 –T133, N216-D79, N216-S78, T237-N60 (2 H-bonds) T256-N60, N96-T199 (2 H-bonds)] (Fig. 4A). Conversely, when the α^43_UTI89^ functional dimer was used as the scaffold, α^43_EDL933^ molecules were predicted to self-associate via one interface encompassing 24 hydrogen bonds [D13-N138 (three hydrogen bonds), T15-N119, G30-T98, T32-N100 (two hydrogen bonds), N60-N96, N60-T98, N60-T80, D79-N60, N60-N60, D79-D79, D79-N60, T80-N60, T98-N60,, T98-G30, N96-N60 N100-T32 (two hydrogen bonds), N119-T15, N138-D13 (three hydrogen bonds)] (SI Appendix Fig. S4). We designed two sets of mutants to discern between Ag43^EDL933^ double or single interface self-association. To test for the single interface, we mutated T15, T32, N60, D79, T98, N100, N119 and N138 to glycine residues (Ag43^EDL933-*mt-single*^). We generated a second mutant where we mutated T199 and T256 to glycine at the base of α^43_EDL-933^ (Ag43^EDL933-*mt-double*^), mapping far from the predicted single interface, and responsible for 10 out of the total predicted 24 hydrogen bonds stabilising the putative Ag43^EDL933^ double interface dimer. Examination of MS528 strains expressing these mutants in cell-aggregation assays showed that while modification of the predicted Ag43^EDL933^ single interface did not alter its aggregative function, mutation of two residues in the double interface completely abolished Ag43^EDL933^ mediated cell aggregation (Fig. 4B). This was not due to differences in the expression of the mutant Ag43^EDL933^ proteins on the cell surface, as confirmed by western blotting analysis of heat-released proteins (Fig. 4B, Inset). Hence we confirmed that α^43_EDL933^ interacts via an extensive double interface. Although both dimers are possible, the double interface dimer results in a 550 Å^2^ buried surface area, compared to 260 Å^2^ for the single interface dimer, suggesting that interactions such as van der Waals contacts are important for dictating the preference for the double interface dimer.

### Oligomeric state of Ag43 homologues in solution

To determine the Ag43 homologue self-association properties in solution, we analysed recombinantly expressed and purified proteins by analytical ultracentrifugation sedimentation velocity and small angle scattering experiments.

Sedimentation-coefficient distribution (c(s)) analysis of α^43a^ (Fig. 5C) and α^43_EDL933^ (Fig. 5D) at 2.2 mg ml^−1^, showed that these two proteins exist primarily as dimeric species with standardised sedimentation coefficients (*s*_20,w_) of 4.7 S and 5.2 S, with lower proportions of a monomeric species at 3.5 S and 3.8 S, respectively. Conversely, at the same concentration α^43_UTI89^ (Fig. 5B) and α^43b^ (Fig. 5A) showed higher proportions of monomeric species at 3.9 S and 2.8 S, respectively. Note that both α^43_UTI89^ and α^43_EDL933^ have two-fold higher extinction coefficients than both Ag43a and Ag43b, and so show correspondingly higher absorbance signals. This data supports the capacity of the Ag43 homologues to form dimers in solution, consistent with their ability to promote aggregation. Importantly, the finding that both α^43a^ and α^43_EDL933^ form higher proportions of dimers compared to α^43b^ and α^43_UTI89^ at the same concentration correlates with their faster aggregation kinetics. Additionally, the frictional ratios (f/f_0_) for these experiments ranged from 1.4–1.8 confirming the extended shape of both the monomers and the trans-dimers.

**Fig. 5.**
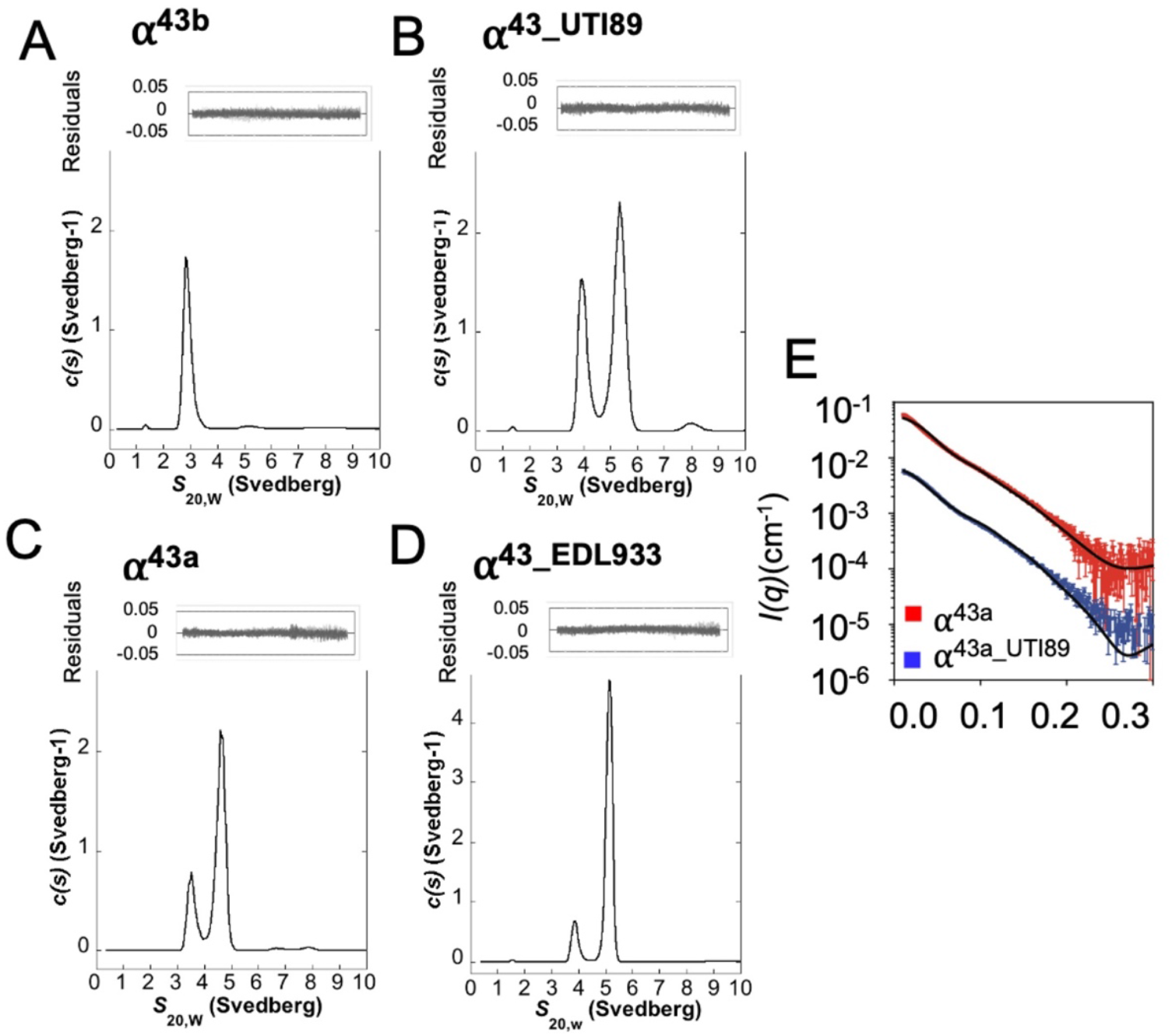
Oligomeric state of Ag43 homologues in solution. Representative analytical ultracentrifugation (AUC) sedimentation velocity analysis of the Ag43 homologues at 2.2 mg/mL with c(s) plotted as a function of *s*_20,w_ (Svedberg). Both α^43a^ *(C)* and α^43_EDL933^ *(D)* showed a much higher proportion of dimer (4.7 and 5.2 S) than monomer (3.5 and 3.8 S). In comparison α^43_UTI89^ (*B*) revealed a higher proportion of monomer (3.9 S) than dimer (5.4 S), whereas α^43b^ *(A)* showed only a monomeric species (2.8 S). Residuals resulting from the *c*(*s*) distribution fits are shown above. Small-angle X-ray scattering measurements of α^43a^ (red) and α^43_UTI89^ (blue) in solution *(E)*. Model scattering curves are overlayed on the experimental data (solid black line) and have been calculated as a linear combination of monomer and dimer scattering curves. The model scattering curves are consistent with the experimental data (*χ*^2^(α^43a^) = 22.9; *χ*^2^(α^43_UTI89^) = 8.8).

We further investigated the solution properties of the two different Ag43 dimer conformations represented by α^43a^ and α^43_UTI89^ using small-angle X-ray scattering (SAXS) (SI Appendix Table S2). In both cases, analysis of the SAXS data found the two Ag43 homologues to be mixtures of different oligomeric states. To understand the way in which these proteins oligomerise, and to quantify the amount of monomer and dimer present in solution, scattering profiles of the monomeric and dimeric forms were calculated from the α^43a^ and α^43_UTI89^ crystal structures, which represent a more extensive double interface and smaller single interface, respectively. The experimental SAXS data were then fit as a linear combination of the model monomer and dimer curves. In both cases, the quality of the fit to the data was good (Fig. 5E), providing evidence that the dimerization occurs in the manner displayed in the crystal structures for both proteins. Further, the analysis indicated that at 1.2 mg mL^−1^, a higher proportion of the protein was present in the dimeric form for α^43a^ compared to α^43_UTI89^, which supports the findings from the analytical ultracentrifugation experiments.

## Discussion

The SAAT subgroup of ATs are major determinants of bacterial aggregation and biofilm formation (9). In an aggregated state, bacteria are more resistant to multiple stresses, including those induced by antibiotics and biocides. The protection offered by bacterial aggregation and biofilm formation is an important contributor to the development of antibiotic resistance and chronic infections, therefore understanding these processes is essential given the ever-increasing threat posed by multidrug-resistant bacteria. While the biology of aggregating autotransporters has been well investigated, understanding the molecular processes underlying bacterial aggregation has been hampered by the lack of atomic detail for the majority of autotransporter proteins that mediate aggregation.

Ag43 is the most prevalent autotransporter in *E. coli* strains and a prototype adhesin for bacterial aggregation studies (9). Phylogenetic network analysis focusing on Ag43 functional domains classified this family of proteins into four major groups (C1-C4) with some groupings showing distinct aggregative properties. (20). In the current study, we have focused on a set of Ag43 variants from three clinically important pathogenic *E. coli* strains EHEC O157:H7 EDL933, UPEC UTI89 and UPEC CFT073. Our characterisation of the cell aggregation phenotypes of Ag43^EDL933^, Ag43^UTI89^ and Ag43b compared to Ag43a, are largely consistent with previous studies (20), but show some further variations. As before, C2 Ag43^EDL933^ shows a strong aggregation phenotype. This feature is also shared by Ag43a from C3, however, this group also displays the highest variability in aggregation kinetics, as it contains other members such as Ag43b which shows a significant delay in autoaggregation. This difference is intriguing considering that these proteins share 85% sequence identity in their functional (α) domains. To our knowledge this is first detailed study of Ag43^UTI89^ bacterial aggregation from C4. Ag43^UTI89^ was found to show a similar delay in aggregation similar to that of Ag43b. These aggregation properties were supported by the differences in bacterial clumping as shown by fluorescence microscopy and flow cytometry with especially Ag43^EDL933^ showing consistently strong aggregation.

To define the mechanistic detail underlying the functional differences observed among the Ag43 variants we sought to determine their crystal structures. Our previous studies on the functional domain of UPEC CFT073 Ag43a (α^43a^), revealed a distinctive curved β-helical architecture and identified that trans-association of Ag43a β-helices from neighbouring cells leads to bacterial clumping (13). In this study, we determined the structure of a further three Ag43 functional domains (α^43_EDL933^, α^43_UTI89^ and α^43b^). Of significance, is the finding that despite sequence identity as low as 45%, all of the tested Ag43 homologues retain the L-shape bend, which is essential for Ag43a bacterial aggregation (13). However, this L-shape is not present in autotransporter adhesins that directly bind host surfaces such as UpaB (43) and pertactin P.69 (32). Thus, it is possible that the L-shape bend may only be required for adhesins that self-associate between bacterial surfaces to promote bacterial aggregation. It remains to be revealed how widespread the L-shape architecture occurs throughout the SAATs and functionally related autotransporters. To date, the only other SAAT structurally determined is TibA from enterotoxigenic *E. coli*, however, this structure only encompasses an N-terminal fragment of the passenger domain (44).

Through structural and mutagenesis studies we have shown that all of the Ag43 homologues examined in this study self-associate in a head-to-tail configuration and that the distinct aggregation dynamics observed for each homologue are determined by their different inter-protein interactions. For example, the rapid aggregation phenotype displayed by Ag43^EDL933^, which mimics that of Ag43a, results from strong α^43_EDL933^-α^43_EDL933^ homotypic interactions which are stabilised by two interfaces with a total of 24 hydrogen bonds (α^43a^ dimers are also stabilised by two interfaces with a total of 18 H-bonds and two salt bridges) (13). Conversely, the delayed aggregation phenotype of Ag43^UTI89^ is a direct result of the weaker α^43_UTI89^ dimers, which are only stabilised by a single interface encompassing 13 hydrogen bonds. The inability of α^43_UTI89^ to form a double interface may be due to the presence of long sidechains on residues such as Arg161 in the F2-F3 loops mapping in the lower part of the β-helix, which in a head-to-tail association via a double interface would result in steric clashes between interacting proteins. Although α^43b^-α^43b^ trans-dimers are stabilised by a double-interface, these only consist of a total of 14 hydrogen bonds, where the lower bonding network results in Ag43b expressing cells displaying a slow aggregation profile comparable to that of Ag43^UTI89^. The differing strengths and configurations of these Ag43 self-associations as interpreted by the crystal structures were confirmed in solution using both AUC and SAXS analysis. Consistent with the aggregation phenotypes both α^43_EDL933^ and α^43a^ showed a higher propensity for self-association compared to α^43_UTI89^ and α^43b^.

Consequently, we reveal how bacterial adhesins such as the Ag43 homologues can acquire simple adaptations to modify their self-association mechanisms. The introduction of additional polar or charged amino acids at their self-association interfaces increases affinity and aggregation kinetics, while acquisition of amino acids opposing in nature perturbs self-association thereby decreases affinity and aggregation kinetics. Interestingly, amongst the Ag43 homologues we investigated they were found to retain a significant degree of conservation (N/D, N, N, X, D, X, N) at their association interfaces. This presumably allows for heterotypic interactions to take place between different Ag43 homologues to influence polymicrobial biofilms between different strains of *E. coli* (20). Of note, α^43_ UTI89^ with the single self-association interface was from the C4 Ag43 homologues that were shown to most readily associate with other Ag43 homologues (20).

Overall, our research reveals that bacterial aggregation is a tightly controlled and complex phenomenon. Apart from regulation of Ag43 at the transcription level by phase variation, the strength of the Ag43 adhesin can also be modified, leading to different degrees of aggregation (Fig. 6). Bacteria are faced with a trade-off, whereby as aggregation is increased, more protection is afforded by the formation of biofilms but bacterial spread is reduced. A main factor which would influence bacterial aggregation and biofilm formation is their environment. UPEC are exposed to varying levels of flushing by urine and host antimicrobial peptides, this is in contrast to EHEC where within the intestine bacteria are exposed to peristalsis and abundant natural microbiota. In some cases, *E. coli* can express more than one Ag43, such as UPEC CFT073 with Ag43a and Ag43b, giving them more versatility (9). The ability of some Ag43 homologues to form hybrid associations with other homologues additionally influences the nature of polymicrobial biofilms. In all, the variations in the phenotypes and mechanisms imparted by different Ag43 homologues are important determinants of pathogen colonisation and adaptation in the human host. Such variations in adhesin attachment are likely to be present in other autotransporter adhesins and even other types of bacterial adhesins, emphasizing the broad importance for these adaptations to different environments.

**Figure 6.**
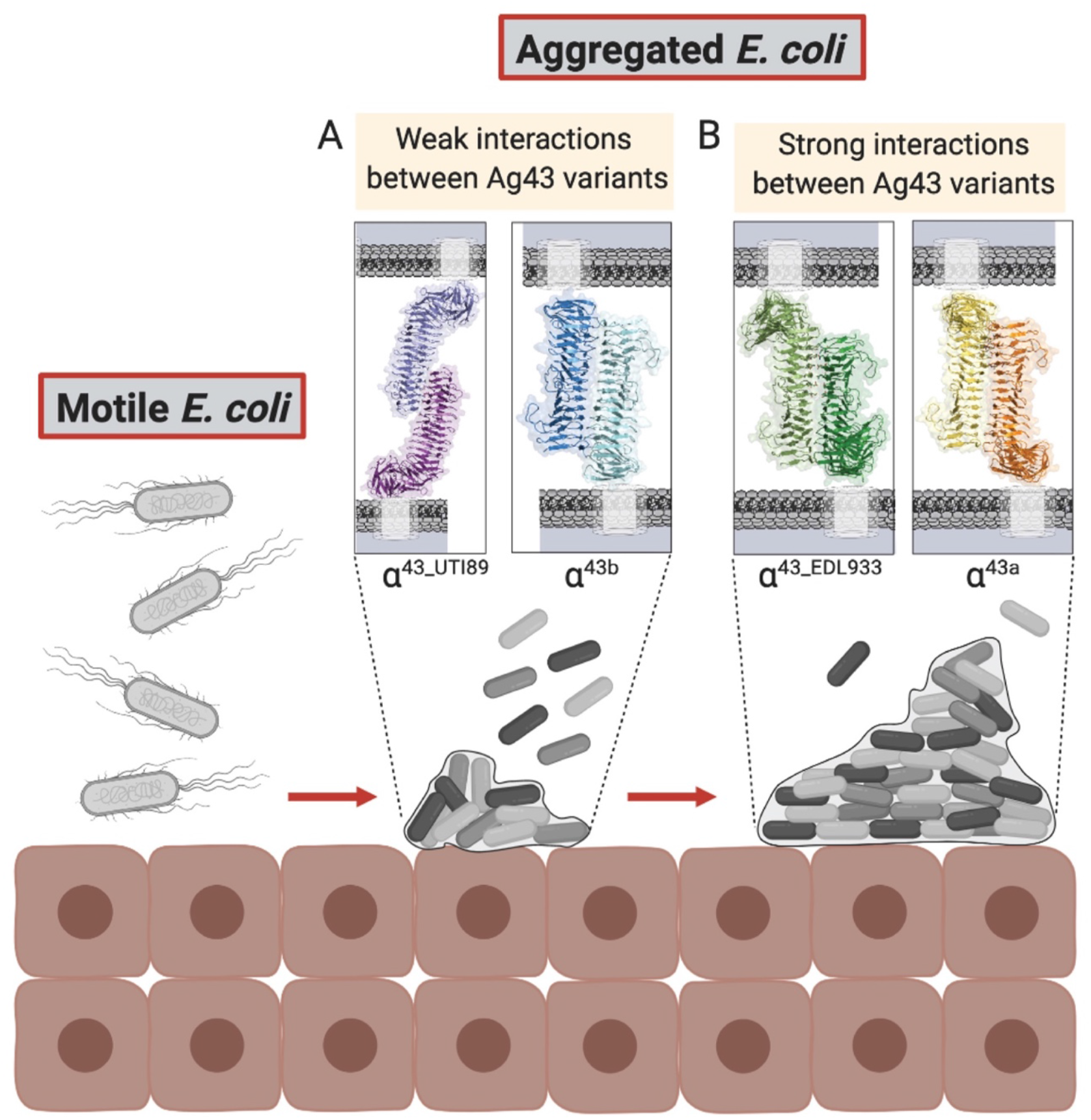
The variations of Ag43 self-association lead to different levels of bacterial aggregation. (*A*) The weaker interactions between Ag43 variants such as Ag43^UTI89^ (13 H-bonds) and Ag43b (14 H-bonds) results in reduced aggregation and biofilm formation and a greater proportion of free bacteria. (*B*) In contrast the higher affinity interactions between Ag43 variants such as Ag43a (18 H-bonds and 2 salt bridges) and Ag43^EDL933^ (24 H-bonds) leads to a more prominent bacterial aggregation and biofilm phenotype. These variations are adaptations by *E. coli* to suit colonisation and persistence in different environments. This image was created with Biorender.

## Acknowledgements

This work was supported by the Australian Research Council (ARC) project grants (DP150102287, DP180102987, DP210100673), Future Fellowship (FT130100580), a National Health and Medical Research Council (NHMRC) Project Grant (GRT1143638) and an NHMRC Fellowship (GRT1106930). Support from INRAE (Institut National de Recherche pour l’Agriculture, l’Alimentation et l’Environnement) and Région Auvergne with FRI IRP (Fond Régional Innovation Institut de Recherche Pharmabiotique) CoMBa grant (AV0003483 and DOS0019690/00) are also acknowledged. We acknowledge the use of the MX1, MX2 and SAXS/WAXS beamlines at the Australian Synchrotron (ANSTO) and the CSIRO Collaborative Crystallisation Centre (www.csiro.au/C3; Melbourne, Australia).

## Materials and Methods

### Cloning of Ag43 homologues

Cloning of *agn43a* (locus tag c3655) and *agn43b* (locus tag c1273) from UPEC CFT073 into pBAD/Myc-His A (45) has been previously described (14). Cloning of *agn43* from UPEC O18:K1:H7 UTI89 (c1139) and *agn43* (*cah*; z1211) from EHEC O157:H7 EDL933 into pBAD/Myc-His A was carried out using primers UTI89_c1139a_Lic_Fw, UTI89_c1139a_Lic_Rv, EDL933_z1211a_Lic_Fw and EDL933_z1211a_Lic_Rv (SI Appendix Table S3). The pBAD/Myc-His plasmids expressing the full length Ag43 proteins were used as the parent vectors for the construction of all mutants, namely Ag43^UTI89-*mt*^, Ag43b^*mt*^, Ag43^EDL933-mt-*single*^, Ag43^EDL933-*mt*-*double*^ (SI Appendix Table S4). All mutant constructs were generated by Epoch Life Science, confirmed by sequencing and used to transform *E. coli* strain MS528 (30, 42) for functional testing.

### Bacterial sedimentation assays

pBAD/Myc-His A plasmids encoding wild-type and mutant versions of Ag43a, Ag43b, Ag43^UTI89^ and Ag43^EDL933^ were transformed into the *E. coli fim agn43* null strain MS528, which is unable to mediate cell aggregation (42). Expression of the proteins was induced by incubation in LB media with 0.2% w/v L-arabinose for 3 h at 37 °C. Bacterial cultures were then adjusted to an optical density at 600 nm (OD_600_) of 2. The cultures were transferred to cuvettes and left to stand at room temperature. OD_600_ measurements were taken at 30 min intervals for 2 h. All assays were performed in triplicate.

### Flow cytometry

pBAD/Myc-His A plasmids encoding Ag43a, Ag43b, Ag43^UTI89^ and Ag43^EDL933^ were transformed into the agn43-negative gfp-positive *E. coli* K-12 strain OS56 (14). Each sample was washed in PBS once and suspended in equal volume of PBS. Sample dilutions in PBS were analysed in triplicate using BD Accuri C6 flow cytometer (BD Bioscience, San Diego, CA, USA) 488 nm laser for excitation and measuring emission at 530/30 nm. Readings were collected in logarithmic mode of at least 5000 events per sample. Data was analysed using FlowJo 10.X.7 (Tree Star).

### Heat release assays and Ag43 immunodetection on the bacterial cell surface

Bacterial cultures were prepared following the same method previously described for sedimentation assays. After adjusting the OD_600_ to 2, the cultures were heated to 60 °C for 2 min to release the α-domains from the cell surface into the supernatant. The heat-released supernatants were TCA precipitated. All samples were analysed by SDS-PAGE and western blot analysis to evaluate the translocation and the production levels of both Ag43 WT and mutant proteins. Rabbit polyclonal serum α^43_UTI89^/ α^43_EDL933^ and α^43a^ were used to detect against α^43_UTI89^, α^43_EDL933^ and α^43b^, respectively.

### Immunofluorescence of bacterial aggregates

*E. coli K-12* strain OS56 harbouring pBAD/Myc-His A plasmids encoding Ag43a, Ag43b, Ag43^UTI89^ and Ag43^EDL933^ were induced by incubation in LB media with 0.2% w/v L-arabinose for 3 h at 37 °C. Each sample was collected at 30 mins for fluorescence microscopy analysis using ZEISS Research Axioplan 2 epifluorescent/light microscope (Carl Zeiss Microimaging). All assays were performed in triplicate.

### Biofilm ring test

The Biofilm Ring Test (BRT) (46) (47) was used to assess biofilm formation at early stages of sessile development of *E. coli fim agn43* null strain MS528 with pBAD/Myc-His A plasmids encoding wild-type Ag43a, Ag43b, Ag43^UTI89^ and Ag43^EDL933^. After adjusting the OD_600_ to 0.01 with sterile LB medium, paramagnetic microbeads were added and homogenised by vortexing prior to loading into 96-well polystyrene microplates and static incubation at 37°C. Control wells were filled with microbeads and LB. For reading at the different time points, wells of microplates were first covered with biofilm control (BFC) contrast liquid prior to scanning before and after 1 min magnetisation using a BFC magnetic rack (BFC, France). Results were expressed as Biofilm Formation Index (BFI); basically, BFI decreases in the course of bacterial sessile development and a BFI ≤ 2 indicates a full immobilisation of the microbeads by bacterial microcolonies (46). All assays were performed in triplicate.

### Expression and purification of Ag43 α-domains

The coding sequences for the functional α-domains of Ag43b (residues 55–444,), Ag43^UTI89^ (residues 53-613) and Ag43^EDL933^ (residues 53-613) were amplified from *E. coli* CFT073, *E. coli* O18:K1:H7 UTI89 and *E. coli* O157:H7 EDL933 respectively using primers shown in SI Appendix Table S3. These sequences were cloned into a modified version of a pMCSG7 expression vector encoding an N-terminal His_6_ tag with thioredoxin (TRX) followed by the tobacco-etch-virus (TEV) protease cleavage site (43). The proteins were expressed in the host strain *E. coli* BL21(DE3)pLysS with the autoinduction method (25 °C for 20 h in the case of Ag43b and 30 °C for 20 h for Ag43^UTI89^ and Ag43^EDL933^). The three proteins were purified by nickel affinity chromatography and TEV-cleaved. Uncleaved proteins were removed by a second nickel affinity chromatography. The α-domains of the proteins were purified to homogeneity by size exclusion chromatography (Superdex 75 GE Healthcare) equilibrated in 25 mM HEPES pH 7, 150 mM NaCl and purity was assessed by sodium dodecyl sulphate-polyacrylamide gel electrophoresis (SDS-PAGE).

### Crystallisation and diffraction data measurement

Purified α^43_UTI89^, α^43_EDL933^ and α^43b^ in 25 mM HEPES, 50 mM NaCl, pH 7 were concentrated to 15.6 mg mL^−1^, 18.3 mg mL^−1^ and 12 mg mL^−1^, respectively, and crystallised using the hanging-drop vapour diffusion method. Small block-like crystals were obtained for α^43_UTI89^ (0.10 × 0.20 × 0.10) from a solution consisting of 20% 2-propanol, 100 mM trisodium citrate/citric acid pH 5.2, 20% PEG 4000. Large needle crystals were obtained for α^43_EDL933^ (0.7 × 0.05 × 0.05) in 100 mM SPG (succinic acid, sodium dihydrogen phosphate and glycine) pH 7.4, 27% PEG 1500. Stacked plate crystals were obtained for α^43b^ (∼0.5 × 0.25 × 0.05) from a solution consisting of 100 mM Na cacodylate pH 6.4, 14% PEG 4000, 20% MPD.

Diffraction data were collected at the Microcrystallography MX2 beamline at the Australian Synchrotron using an ADSC Q315r CCD detector. 180° images were collected for crystals of the 3 proteins, at an oscillation angle of 1°, exposure time of 1 sec. iMosflm (48) and Aimless (49) software was utilised to index, integrate and scale the collected diffraction data.

### Structure determination of Ag43 homologues and refinement

The structure of α^43_UTI89^ and α^43b^ was solved with Phaser (50) by molecular replacement using the structure of α^43a^ (PBD: 4KH3) (13) as a model. Similarly, the structure of α^43_EDL933^ was solved via molecular replacement using α^43_UTI89^ as a reference. The corresponding protein models were manually built in Coot (51), and refined with phenix.refine (52) and TLS (Translation/Libration/Screw) refinement (53). Model quality was monitored by the R-free value, which represented 5% of the data. The models for all homologues were evaluated by MolProbity (54) and figures were generated using PyMOL (51, 55). Coordinates and structure-factor files for α^43_UTI89^, α^43b^ and α^43_EDL933^ have been deposited in the Protein Data Bank, with accession codes 7KO9, 7KOB, 7KOH respectively. Data-processing and refinement statistics are summarised in Table 1.

### Analytical ultracentrifugation

Sedimentation velocity experiments were performed using a Beckman Optima XL-A analytical ultracentrifuge, 8-hole An-50 Ti rotor. Protein samples (380 µL) obtained after size exclusion chromatography of the recombinant Ag43 homologues in 25mM HEPES, 150mM NaCl, pH 7.0 and reference buffer (400 µL) were loaded into double-sector quartz cells. Initial scans were performed at 725 × g to determine the optimal wavelength and radial positions. Absorbance readings were collected at 285 nm and 128,794 × g at 20 °C. Solvent density, solvent viscosity and estimates of the partial specific volume of α^43b^ (0.7211 mL g^−1^), α^43_UTI89^ (0.7197 mL g^−1^), α^43a^ (0.7215 mL g^−1^) and α^43_EDL933^ (0.7207 mL g^−1^) at 20 °C were calculated with SEDNTERP (56). Data were analysed using c(s) with SEDFIT (57).

### Small Angle X-ray Scattering

SAXS data were collected on the SAXS/WAXS beamline at the Australian Synchrotron (58). Approximately 60 μL of α^43a^ and α^43_UTI89^, at 1.2 mg mL^−1^, in 25 mM HEPES, 150 mM NaCl, pH 7.4 were loaded into a 1 mm quartz capillary. Samples were flowed during data collection to reduce radiation damage. Data reduction was carried out using the ScatterBrain software (v.2.71) and the data were corrected for solvent scattering and sample transmission, then radially averaged to produce I(*q*) as a function of *q*, where *q* = (4*π*sin*θ*)/*λ, θ* is half the scattering angle, and *λ* is the X-ray wavelength. For α^43a^ (SASBDB ID: SASDKQ3) and α^43_UTI89^ (SASBDB ID: SASDKP3), data were fit as a linear combination of model curves to quantify the nature and proportions of different oligomeric states in solution (59). Model scattering curves were calculated using CRYSOL (v.2.8.3) (60), using monomer and dimer structures taken from PDB: 4KH3 for α^43a^ and PDB: 7KO9 for α^43_UTI89^.

## Data availability

The crystallography, atomic coordinates, and structure factors reported in this paper have been deposited in the Protein Data Bank, www.pdb.org (PDB ID codes 7KO9, 7KOB and 7KOH). The Small Angle X-ray Scattering data reported in this paper have been deposited in the Small Angle X-ray Scattering Biological Data Bank, www.sasbdb.org (SASBDB ID codes: SASDKP3 and SASDKQ3).

## Author Contributions

B.H. and M.A.S designed the research. J.L.V, G.C.M.O, M.T, A.W.L, A.E.W, L.H, K.M.P, V.A and N.C performed the experiments. A.W.L, A.E.W, M.D, M.A.S, J.J.P and B.H analysed the data. B.H, J.J.P, M.D and M.A.S supervised aspects of the work. All authors contributed to the interpretation of the results. B.H, J.J.P and M.A.S. wrote the manuscript. All authors contributed to the critical revision of the manuscript, read and approved the final manuscript.

## Supplementary Information

**Fig. S1.**
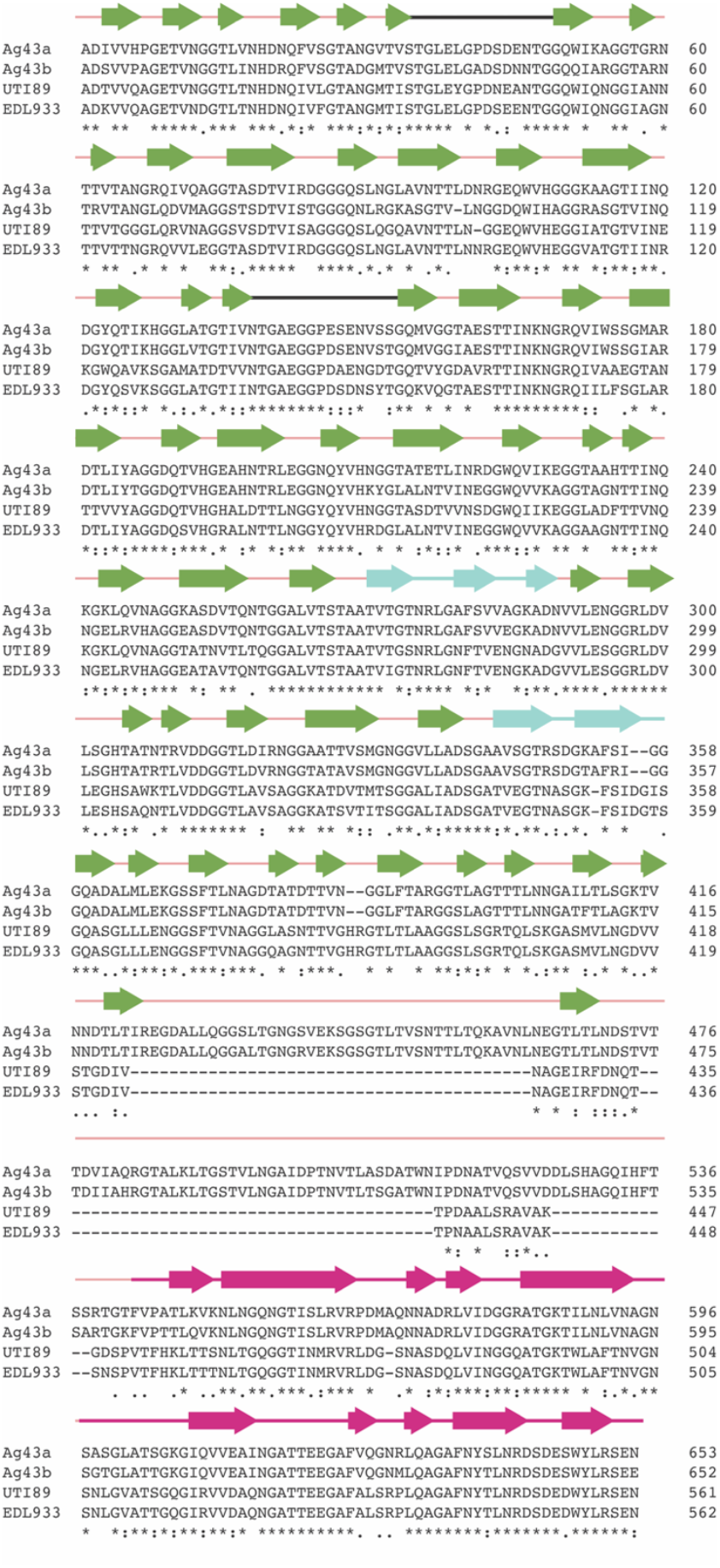
Multiple sequence alignment of α^43a^, α^43b^, α^43_EDL933^ and α^43_UTI89^. (*A*) Sequence alignment of the α-domains of Ag43a, Ag43b, Ag43^UTI89^ and Ag43^EDL933^ obtained with Clustal Omega. Secondary structural elements are based on the structure of α^43_EDL933^, following the colour coding of Fig 2A: β-strands illustrated in green and loops depicted in salmon. The two protruding loops are shown in black, the β-hairpins appear in cyan and the region coloured in hot pink corresponds to the AC domain.

**Fig. S2.**
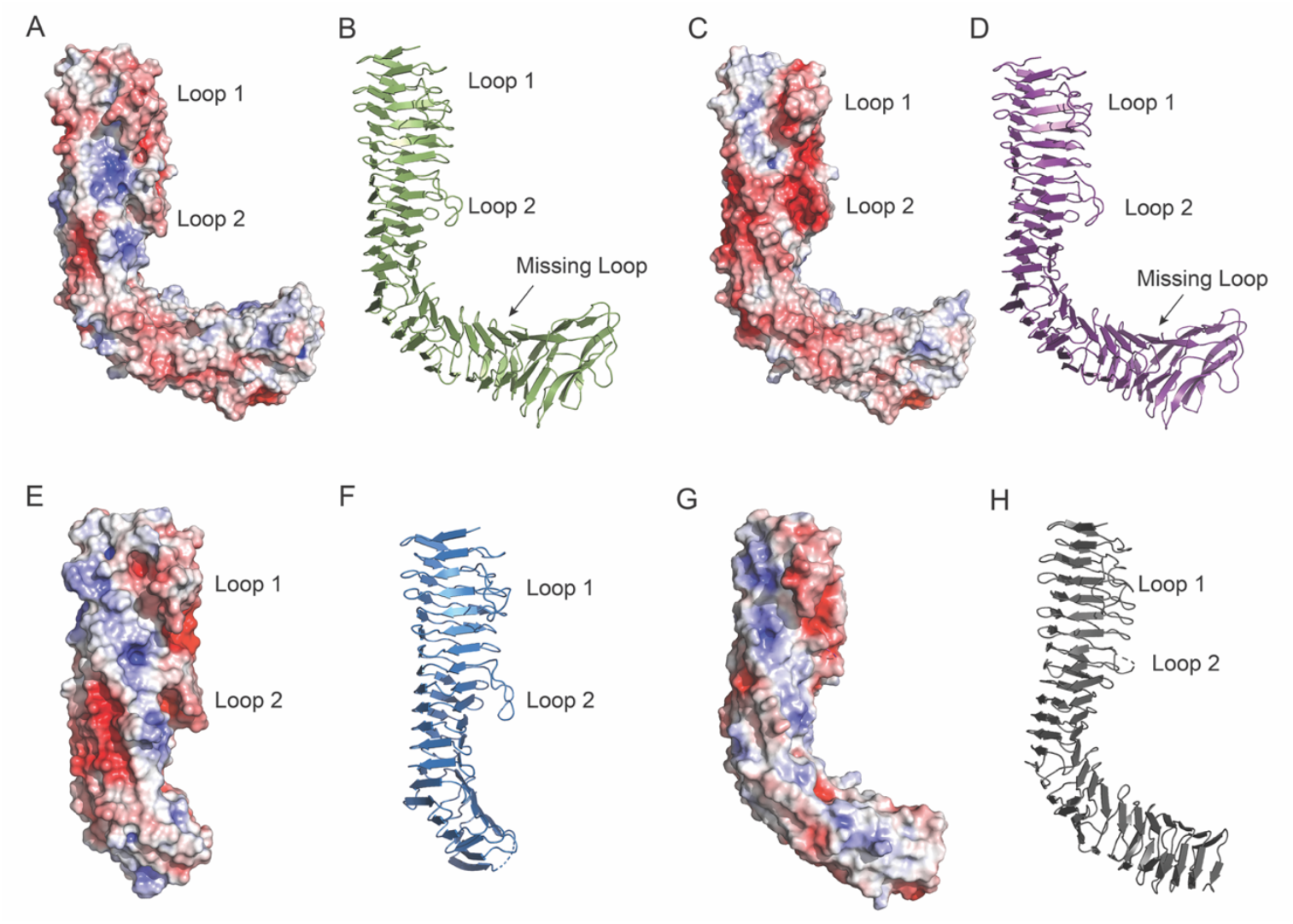
Electrostatic surface representation of α^43_EDL933^, α^43_UTI89^, α^43b^ and α^43a^. (*A, C, E, G*) Electrostatic surface representation of α^43_EDL933^, α^43_UTI89^ and α^43b^ respectively, along with the previously published α^43a^ (PDB: 4HK3) for comparison. For each protein, positive electrostatic potentials are shown in blue, while negative electrostatic potentials appear in red (saturation at 5 kT/e). The two loops (Loop 1 and Loop 2) that protrude from the β-helices reveal acidic patches in these negatively charged loops. (*B, D, F, H*) Cartoon representation of α^43_EDL933^, α^43_UTI89^, α^43b^ and α^43a^ respectively showing the orientation of all proteins in panels A, C, E and G. Figures of the electrostatic potentials were generated using APBS (1).

**Fig. S3.**
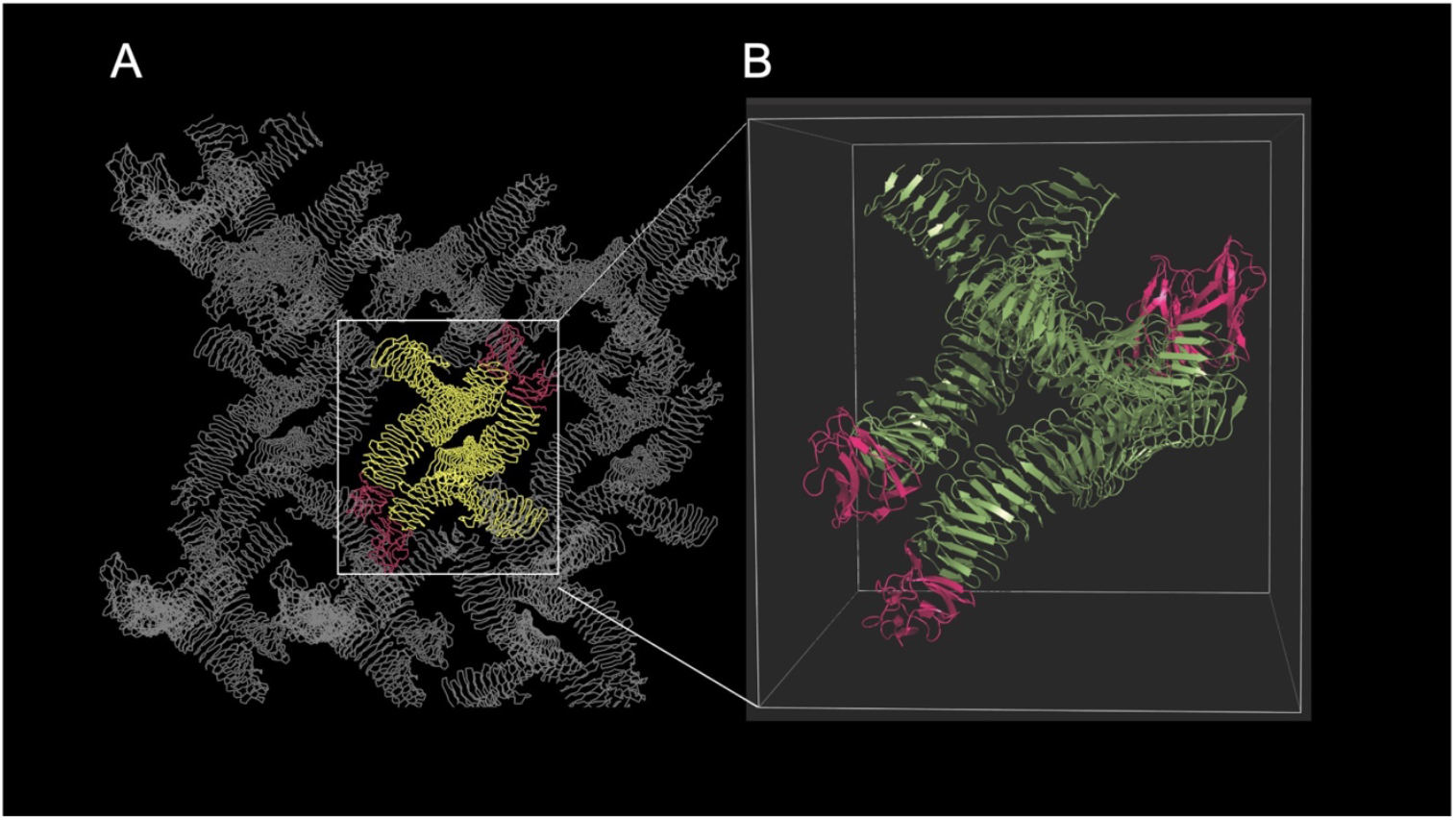
(*A*) Crystal lattice of α^43_EDL933^ with non-biologically relevant crystal contacts observed between the molecules. (*B*) A zoomed view of α^43_EDL933^, showing the four molecules present in its asymmetric unit. The α-domains are depicted in green and the AC domains are displayed in hot pink.

**Fig. S4.**
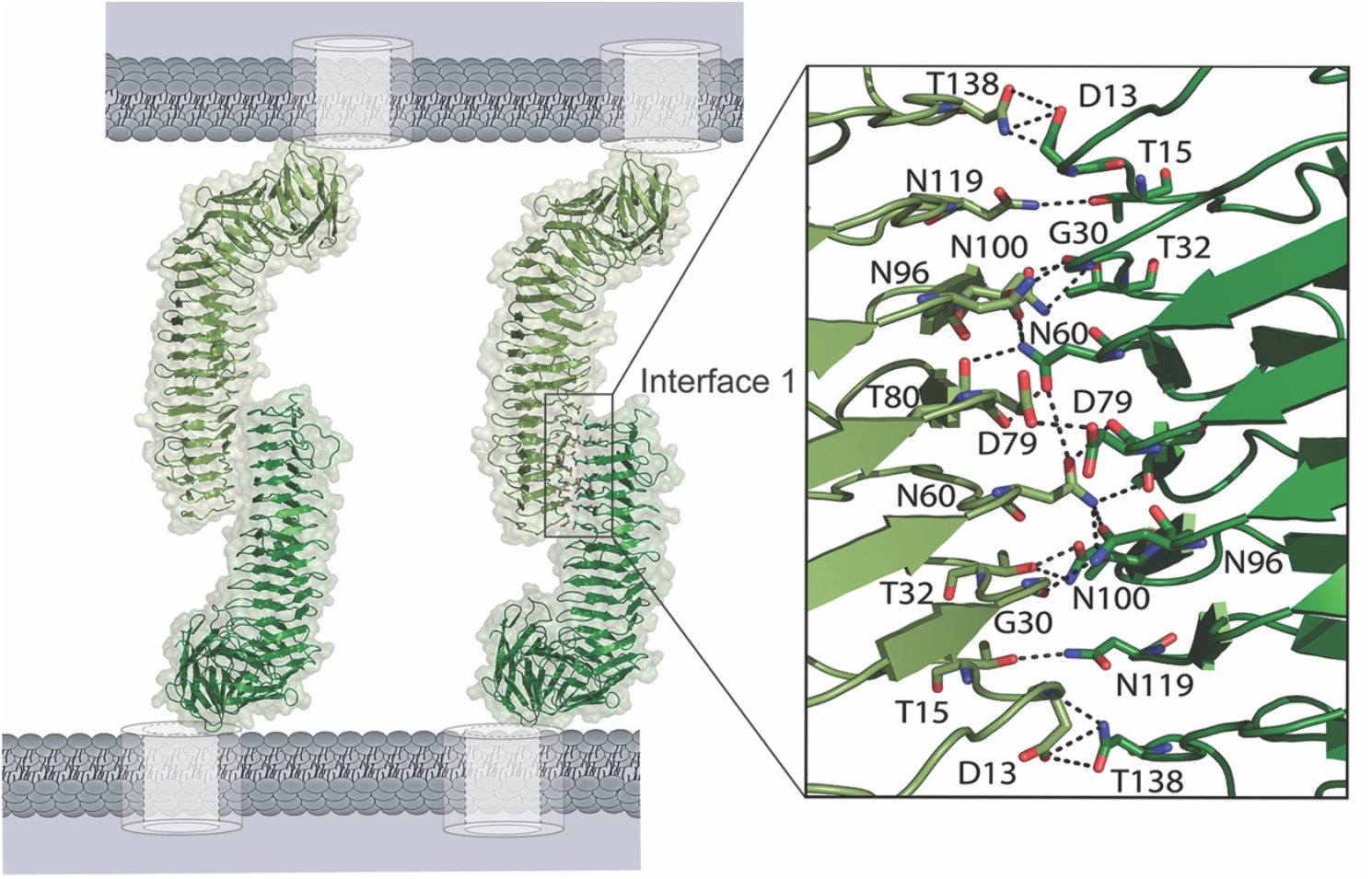
α^43_EDL933^-α^43_EDL933^ dimeric single-interface. Self-association of α^43_EDL933^ molecules on the bacterial cell surface, showing a single interface predicted using α^43_UTI89^ dimer as a model. A close-up view of the interaction interface is shown; the interface consists of 24 hydrogen bonds [D13-N138 (three hydrogen bonds), T15-N119, G30-T98, T32-N100 (two hydrogen bonds), N60-N96, N60-T98, N60-T80, D79-N60, N60-N60, D79-D79, D79-N60, T80-N60, T98-N60, T98-G30, N96-N60 N100-T32 (two hydrogen bonds), N119-T15, N138-D13 (three hydrogen bonds)]. This dimer forms through self-interaction via the F3 face of α^43_EDL933^.

**Table S1.** Comparison of the structures of Ag43 passenger (α) domains and Ag43 autochaperone (AC) domains with reference autotransporter AC domains.

| | $\alpha^{43a}$<br>(492)<sup>d</sup> | $\alpha^{43\_UTI89}$<br>(540)<sup>d</sup> | $\alpha^{43\_EDL933}$<br>(545)<sup>d</sup> | $\alpha^{43b}$<br>(361)<sup>d</sup> | $\alpha^{43\_EDL933-AC}$<br>(109)<sup>d</sup> | $\alpha^{43\_UTI89-AC}$<br>(109)<sup>d</sup> | $\alpha^{EspP-AC}$<br>(131)<sup>d</sup> | $\alpha^{Hap-AC}$<br>(124)<sup>d</sup> | $\alpha^{Hbp-AC}$<br>(97)<sup>d</sup> | $\alpha^{IcsA-AC}$<br>(135)<sup>d</sup> | $\alpha^{P69-AC}$<br>(96)<sup>d</sup> |
| --- | --- | --- | --- | --- | --- | --- | --- | --- | --- | --- | --- |
| $\alpha^{43\_EDL933}$<br>(545)<sup>d</sup> | 2.09<sup>a</sup><br>(413)<sup>b</sup><br>(64.6%)<sup>c</sup> | 2.07<sup>a</sup><br>(493)<sup>b</sup><br>(74.6%)<sup>c</sup> | 0<sup>a</sup><br>(545)<sup>b</sup><br>(100%)<sup>c</sup> | 0.58<sup>a</sup><br>(354)<sup>b</sup><br>(71.2%)<sup>c</sup> | | | | | | | |
| $\alpha^{43\_UTI89}$<br>(540)<sup>d</sup> | 1.62<sup>a</sup><br>(455)<sup>b</sup><br>(57.6%)<sup>c</sup> | 0<sup>a</sup><br>(540)<sup>b</sup><br>(100%)<sup>c</sup> | 2.07<sup>a</sup><br>(493)<sup>b</sup><br>(74.6%)<sup>c</sup> | 2.54<sup>a</sup><br>(306)<sup>b</sup><br>(50.3%)<sup>c</sup> | | | | | | | |
| $\alpha^{43b}$<br>(361)<sup>d</sup> | 0.52<sup>a</sup><br>(350)<sup>b</sup><br>(80.9%)<sup>c</sup> | 2.54<sup>a</sup><br>(306)<sup>b</sup><br>(50.3%)<sup>c</sup> | 0.58<sup>a</sup><br>(354)<sup>b</sup><br>(71.2%)<sup>c</sup> | 0<sup>a</sup><br>(361)<sup>b</sup><br>(100%)<sup>c</sup> | | | | | | | |
| $\alpha^{43\_EDL933-AC}$<br>(109)<sup>d</sup> | | | | | 0<sup>a</sup><br>(109)<sup>b</sup><br>(100%)<sup>c</sup> | 0.38<sup>a</sup><br>(109)<sup>b</sup><br>(98.2%)<sup>c</sup> | 2.20<sup>a</sup><br>(91)<sup>b</sup><br>(23.1%)<sup>c</sup> | 2.96<sup>a</sup><br>(76)<sup>b</sup><br>(26.3%)<sup>c</sup> | 4.36<sup>a</sup><br>(21)<sup>b</sup><br>(19%)<sup>c</sup> | 2.53<sup>a</sup><br>(92)<sup>b</sup><br>(27.2%)<sup>c</sup> | 2.79<sup>a</sup><br>(69)<sup>b</sup><br>(16%)<sup>c</sup> |
| $\alpha^{43\_UTI89-AC}$<br>(109)<sup>d</sup> | | | | | 0.38<sup>a</sup><br>(109)<sup>b</sup><br>(98.2%)<sup>c</sup> | 0<sup>a</sup><br>(109)<sup>b</sup><br>(100%)<sup>c</sup> | 3.50<sup>a</sup><br>(80)<sup>b</sup><br>(7.5%)<sup>c</sup> | 2.83<sup>a</sup><br>(76)<sup>b</sup><br>(26.3%)<sup>c</sup> | 2.52<sup>a</sup><br>(14)<sup>b</sup><br>(14.3%)<sup>c</sup> | 2.56<sup>a</sup><br>(92)<sup>b</sup><br>(27.2%)<sup>c</sup> | 2.28<sup>a</sup><br>(58)<sup>b</sup><br>(19%)<sup>c</sup> |

7KO9, 7KOB, 7KOH
<sup>a</sup>RMSD (Root Mean Square Deviation) values (Å) calculated using Secondary Structure Matching (SSM) superimpose tool in Coot (2)
<sup>b</sup> number of aligned Cα atoms
<sup>c</sup> sequence identity
<sup>d</sup> total number of residues
Aligned structures: Ag43a (PDB: 4KH3), Ag43<sup>UTI89</sup> (PDB: 7KO9), Ag43<sup>EDL933</sup> (PDB: 7KOH), Ag43b (PDB: 7KOB), Ag43<sup>EDL933</sup> (PDB: 7KOH; AC: V<sub>453</sub>-E<sub>561</sub>), Ag43<sup>UTI89</sup> (PDB: 7KO9; AC: V<sub>452</sub>-E<sub>560</sub>), EspP (PDB: 3SZE; AC: D<sub>869</sub>-A<sub>999</sub>), Hap (PDB: 3SYJ; AC: D<sub>830</sub>-P<sub>976</sub>), Hbp (PDB: 1WXR; AC: V<sub>946</sub>-N<sub>1048</sub>), P69 (PDB: 1DAB; AC: L<sub>444</sub>-P<sub>539</sub>) and IcsA (PDB: 3ML3; AC: D<sub>606</sub>-D<sub>740</sub>)

**Table S2.** SAXS data collection details.

| | $\alpha^{43a}$ | $\alpha^{43\_UTI89}$ |
| --- | --- | --- |
| SASBDB ID | SASDKQ3 | SASDKP3 |
| Data Collection Parameters |  |  |
| Instrument | SAXS-WAXS, Australian Synchrotron | SAXS-WAXS, Australian Synchrotron |
| Beam geometry ($\mu\text{m}$) | $80 \times 200$ | $80 \times 200$ |
| Wavelength ($\text{\AA}$) | 1.0332 | 1.0332 |
| Flux (photons/s) | $3.1 \times 10^{12}$ | $3.1 \times 10^{12}$ |
| Sample to detector distance (m) | 2.680 | 1.428 |
| $q$-range ($\text{\AA}^{-1}$) | 0.00–0.30 | 0.011–0.60 |
| Temperature (K) | 285 | 285 |
| Absolute intensity calibration | Water | Water |
| Exposure time (s) | 14 ($14 \times 1$ s exposures) | 35 ($35 \times 1$ s exposures) |
| Configuration | Single measurement from 96-well plate | Single measurement from 96-well plate |
| Protein concentration ($\text{mg ml}^{-1}$) | 1.2 | 1.2 |

**Table S3.**
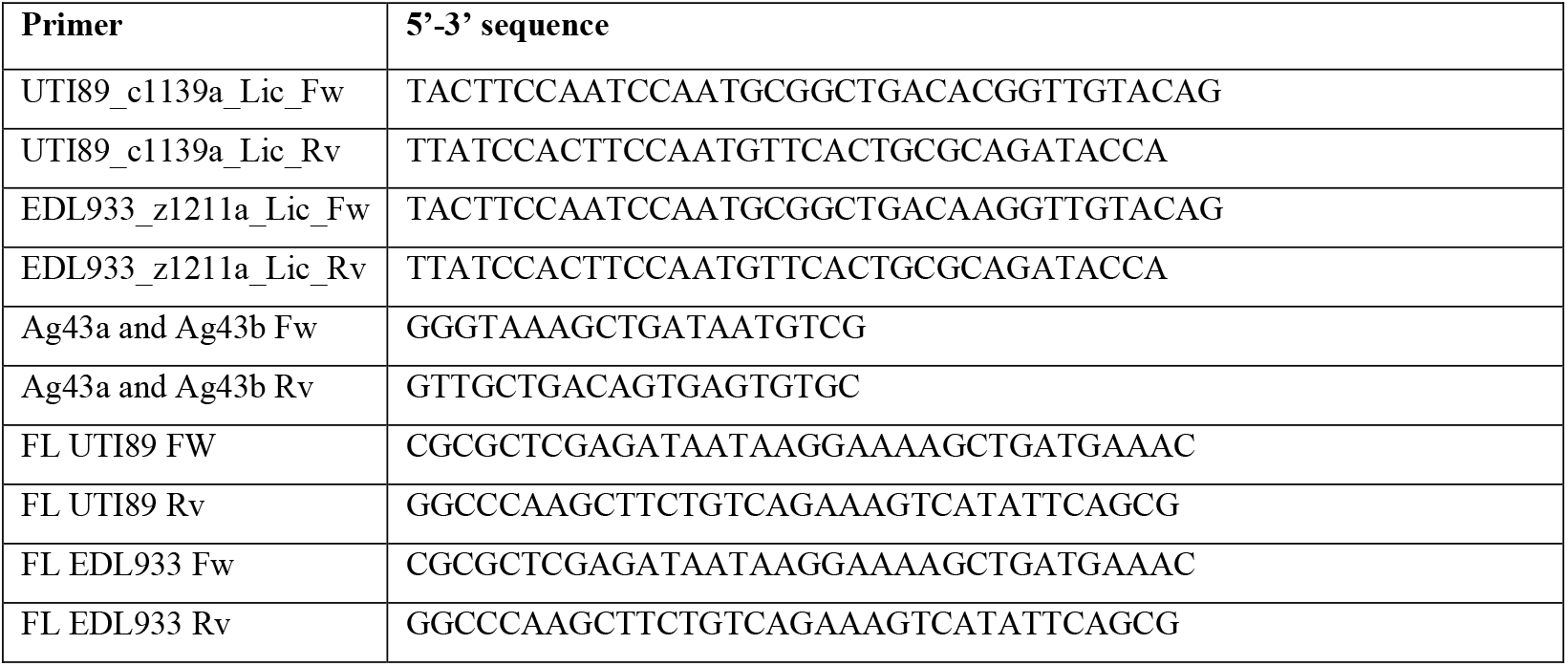
Primers used in this study.

**Table S4.**
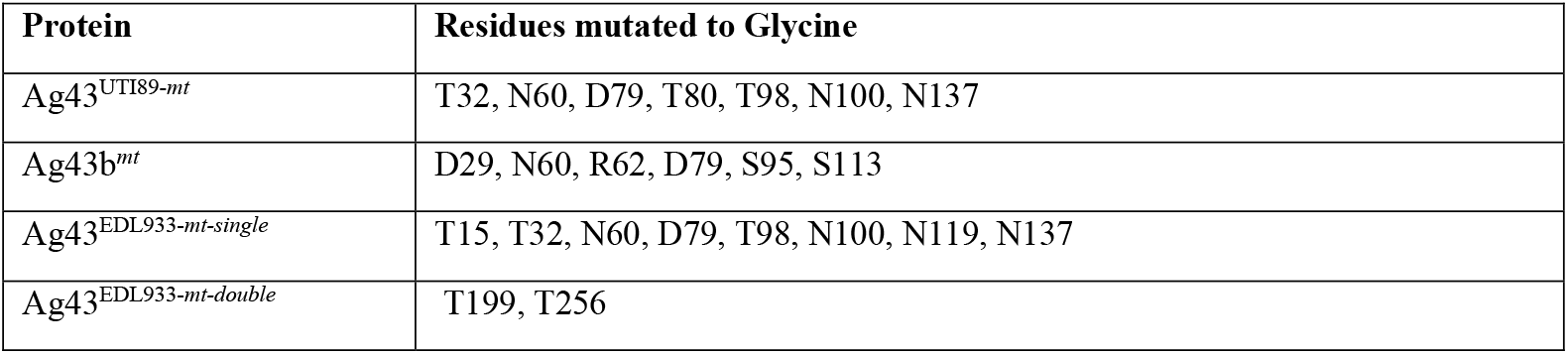
Residues mutated for mutant design of proteins.

